# Genomics implicates adaptive and innate immunity in Alzheimer’s and Parkinson’s

**DOI:** 10.1101/059519

**Authors:** Sarah A Gagliano, Jennie G Pouget, John Hardy, Jo Knight, Michael R Barnes, Mina Ryten, Michael E Weale

## Abstract

**Objectives:** We assessed the current genetic evidence for the involvement of various cell types and tissue types in the aetiology of neurodegenerative diseases, especially in relation to the neuroinflammatory hypothesis of neurodegenerative diseases.

**Methods:** We obtained large-scale genome-wide association study (GWAS) summary statistics from Parkinson’s disease (PD), Alzheimer’s disease (AD), and amyotrophic lateral sclerosis (ALS). We used multiple sclerosis (MS), an autoimmune disease of the central nervous system, as a positive control. We applied stratified LD score regression to determine if functional marks for cell type and tissue activity, and gene set lists were enriched for genetic heritability. We compared our results to those from two gene-set enrichment methods (Ingenuity Pathway Analysis and enrichr).

**Results:** There were no significant heritability enrichments for annotations marking genes active within brain regions, but there were for annotations marking genes active within cell-types that form part of both the innate and adaptive immune systems. We found this for MS (as expected) and also for AD and PD. The strongest signals were from the adaptive immune system (e.g. T cells) for PD, and from both the adaptive (e.g. T cells) and innate (e.g. CD14: a marker for monocytes, and CD15: a marker for neutrophils) immune systems for AD. Annotations from the liver were also significant for AD. Pathway analysis provided complementary results.

**Interpretation:** For Alzheimer’s and Parkinson’s disease, we found significant enrichment of heritability in annotations marking gene activity in immune cells.

## Introduction

Neurodegenerative diseases – including Alzheimer’s (AD), amyotrophic lateral sclerosis (ALS), and Parkinson’s disease (PD) – are personally devastating and an increasing burden on health-care systems worldwide. Recently there has been much progress in identifying genetic variants associated with neurodegenerative diseases. In the latest AD meta-analysis 19 loci in addition to the well-established APOE locus were pinpointed.^1^ The latest ALS meta-analysis identified three ALS-associated loci^2^ and the latest PD meta-analysis brought the total number of established PD loci to 26.^3^ Despite progress in identifying genetic hits in these neurodegenerative diseases, the underlying processes or cell-types mediating the pathology remain uncertain.

As genome-wide association studies (GWASs) have grown in size and power, so has the quality and scope of functional information that can be used to annotate the genome with relevant genomic and epigenomic marks linked to the regulation of gene expression. Previous studies have demonstrated enrichment of disease-associated variants (for numerous diseases) with functional genomic annotations, including DNase I hypersensitive sites, transcription factor binding sites, histone modifications, and expression quantitative trait loci (eQTLs).^4–7^ These annotations vary depending on cell/tissue-type. Given the many ways in which complex diseases arise, and for human brain diseases, the well-recognized cellular heterogeneity of the brain, pinpointing cell-types of interest is important to further understand pathogenicity. Efforts to obtain brain samples (the most obviously relevant tissue for neurodegenerative diseases) for eQTL analyses are ongoing.^8–13^ There has been a recent proliferation in the availability of cell-type and tissue-specific annotations, including brain tissue, for example through the Roadmap Epigenomics Project^14^ and the PsychEncode Project.^11^ Nevertheless, obtaining large numbers of post-mortem human brains remains challenging, and current eQTL analyses are likely to be underpowered. Characterization of eQTLs and DNA regulatory elements in blood is a complementary approach.

The neuroinflammatory hypothesis of neurodegenerative diseases posits that dysregulation of the immune system is an important factor in the aetiology of these diseases.^15,16^ There is little doubt that multiple sclerosis (MS) is an immune-mediated disease^17 18,19^ We therefore use this disease as a positive control with regard to expected enrichment in heritability for annotations from immune cells. There is extensive functional and clinical evidence that immune dysfunction plays a key role in the pathogenesis of the relapse-remitting phase of MS.^20,21^ For AD, Yokoyama *et al*.^22^ showed that eight variants were associated with both AD and immune-mediated diseases, and there is further evidence from pathway analysis^1,23,24^ and from animal models.^25^ For PD, the role of the immune system has been suggested through pathway analysis^26,27^, animal models^28^, and variants in the HLA region reaching statistical significance in genome-wide association studies.^3,29^ For ALS, there is evidence of immune abnormalities.^30^ Nevertheless, the extent to which the immune system is involved in neurodegenerative diseases such as AD, ALS and PD, and the potential roles played by the innate and adaptive immune components, remain unestablished.

Finucane *et al*.^31^ introduced stratified LD score regression as a method for partitioning the inferred heritability from GWAS summary statistics. They determined whether genetic heritability in 17 GWASs was enriched within various functional annotations which reflected parts of the genome that were active in a number of tissues and cell-types. We applied this methodology to four diseases (MS, AD, ALS, PD) to test for enrichment of heritability, both using Finucane *et al.*’s^31^ cell-type group annotations and using additional annotations from brain and immune cells and from published sets of brain and immune-related genes.^32^

## Methods

We obtained GWAS summary statistics for three neurodegenerative diseases: AD,^1^ ALS^2^ and PD.^3^ We used MS^33^ as a positive control, as it is a disease affecting the brain with known immune aetiology. All studies were conducted in European populations, and are summarized in Table 1. For AD, which is a two-stage study, we only used data from the first stage. (See Box 1 for details on this study.) We did not study Huntington’s disease (which has other genetic modifiers in addition to the primary *HTT* locus) and frontotemporal dementia, because the current GWAS sample sizes for these diseases are modest, and thus the datasets were considered to be insufficiently powered for our analyses.^31^

**Table 1.** Description of the GWASs summary statistics

| Neurodegenerative disease | PMID | Cases | Controls | Cohorts |
| --- | --- | --- | --- | --- |
| Parkinson's disease (PD) | 25064009 | 13,708 | 95,282 | 15 |
| Alzheimer's disease (AD) | 24162737 | 17,008 | 37,154 | 19 |
| Amyotrophic lateral sclerosis (ALS) | 24256812 | 7,177 | 8,393 | 8 |
| Multiple sclerosis (MS) | 21833088 | 9,772 | 17,376 | 23 |

### Box 1.

International Genomics of Alzheimer’s Project (IGAP) is a large two-stage study based upon genome-wide association studies (GWAS) on individuals of European ancestry. In stage 1, IGAP used genotyped and imputed data on 7,055,881 single nucleotide polymorphisms (SNPs) to meta-analyse four previously-published GWAS datasets consisting of 17,008 Alzheimer’s disease cases and 37,154 controls (The European Alzheimer’s disease Initiative – EADI the Alzheimer Disease Genetics Consortium – ADGC The Cohorts for Heart and Aging Research in Genomic Epidemiology consortium – CHARGE The Genetic and Environmental Risk in AD consortium – GERAD). In stage 2, 11,632 SNPs were genotyped and tested for association in an independent set of 8,572 Alzheimer’s disease cases and 11,312 controls. Finally, a meta-analysis was performed combining results from stages 1 & 2.

We estimated pairwise genetic correlations among the four diseases using cross-trait LD score regression.^34^ We then applied stratified LD score regression to determine if various functional categories (cell-type groups, annotations at the tissue/cell level for brain or immune cells, and sets of brain and immune gene lists) were enriched for heritability. LD score regression exploits the expected relationships between true association signals and local LD around them to correct out systematic biases and arrive at unbiased estimates of genetic heritability within a given set of SNPs (here stratified according to their functional category).^31^ Following Finucane *et al*.^31^, we added annotations individually to the baseline model; we used HapMap Project Phase 3 SNPs for the regression and 1000 Genomes Project European population SNPs for the reference panel; we only partitioned the heritability of SNPs with minor allele frequency >5%; and we excluded the MHC region from analysis. The high LD and strong association signals within the MHC region result in a dominating effect on LD score regression, and for the purposes of our analyses excluding this region results in a conservative approach.

The grouped cell-type annotations provided by Finucane *et al*.^31^ are the union of histone marks for 10 broad categories including central nervous system (CNS), cardiovascular, immune/hematopoietic, and liver. For these analyses we corrected for multiple testing of four GWASs across 10 cell-type groups (4 × 10 = 40 hypotheses tested), resulting in a Bonferroni significance threshold of p= 1.2 × 10^−3^.

We then extended the analytical approach of Finucane *et al.*^31^ in the following ways. Firstly, we obtained additional annotation information. We obtained histone marks and DNase I hypersensitive sites data from the Roadmap Epigenomics Consortium;^14^ we obtained eQTLs derived from brain regions from the UK Brain Expression Consortium^10^ and the GTEx Consortium;^9^ and we obtained promoter capture HiC array express data in CD34 (a marker of immature hematopoietic cells) cells from GM12878 (reference: E-MTAB-2323).^35^ We also considered two gene-sets. All these annotations are listed in **Supplementary Table 1**. Secondly, in order to reduce the multiple testing burden, we combined information across the four different histone marks and the DNase I hypersensitive marks in order to create an aggregate set of regulatory marks for each cell type. This aggregation annotation was obtained based on a simple union operation: for each tissue or cell-type, a SNP was labeled as ‘annotated’ based on whether it possessed any relevant histone mark or DNase I hypersensitive mark. Both DNase I and histone marks are known to reflect active regions of the genome, motivating their aggregation in order to create a general mark of genomic activity. DNase I sites are associated with an open chromatin structure, and different histone marks are markers of active promoters (H3K4Me3 + H3K27Ac) or active enhancers (H3K4Me1 + H3K27Ac) regions.

For brain tissue, we defined a union set of histone marks plus DNase I hypersensitive sites from the Roadmap Epigenomics Consortium^14^ using the same aggregation procedure as above. This processing resulted in one annotation per brain region (10 annotations).

We grouped eQTLs across all brain regions, but treated the eQTLs from the UK Brain Expression Consortium^10^ and the GTEx Consortium^9^ separately (resulting in two annotations). Both the GTEx and UKBEC analyses included brain regions highly relevant to MS, AD and PD, namely white matter, hippocampus, temporal cortex and substantia nigra.

Amongst specific immune cells we assessed the histone marks previously described,^31^ and also the histone marks and DNase I hypersensitive site data from the Roadmap Epigenomics Consortium for immune and blood cells,^14^ and we took the union for each cell-type as described above (three histone marks and DNase I hypersensitive site). This resulted in 20 annotations from Finucane *et al*.^31^ and 14 annotations from Roadmap.

Additionally, we defined four immune cell-type annotations based on promoter capture HiC array express data in CD34 from GM12878 (reference: E-MTAB-2323).^35^ The data for the prey and bait were analysed separately for interactions between captured promoter and captured promoter interactions and for captured promoter and all other regions, which resulted in four annotations.

The above cell/tissue-type specific annotations resulted in a multiple testing correction for four GWASs across 50 (12 brain + 38 immune) annotations (4 × 50 = 204 hypotheses tested). Thus we set a Bonferroni significance threshold of p= 2.5 × 10^−4^ for these analyses. Note that there are correlations within the immune and brain annotations, making our Bonferroni correction somewhat conservative

We also applied heritability enrichment analysis to two sets of genes; one with known brain and one with known immune function. We used a brain gene list of 2,635 genes previously described by Raychaudhuri *et al*.^36^, and an immune gene list of 973 genes previously described by Pouget *et al*.^32^ Brain genes were defined as those fulfilling any of the following criteria: preferential expression in the brain compared to other tissues, “neural-activity” annotation in Panther, “learning” annotation in Ingenuity, and “synapse” annotation in Gene Ontology. Immune genes were defined as those with an “immune response” annotation in at least three of the following databases: Kyoto Encyclopedia of Genes and Genomes, Gene Ontology, Ingenuity, and Immunology Database and Analysis Portal. SNPs were annotated to genes using a 50 kb window, and a baseline list of all genes using this 50 kb window was included in the model as previously described.^32^

Finally, we contrasted the above heritability enrichment analyses with a complementary approach based on gene-set enrichment analysis. We used Ingenuity Pathway Analysis (IPA) (www.ingenuity.com) to identify pathway enrichment among genes associated with different neurological traits for canonical pathways. Canonical pathways are structured pathways. Data from the different phenotypes were integrated and subjected to network analysis via IPA to identify pathway enrichment.Enriched networks are ordered by −log p-value, based on a Fisher Exact test p-value.^37^ For each disease we included SNPs with a p-value <5×10^−4^ and excluded SNPs in the MHC region due to the long stretches of LD in this region. We also performed a pathway analysis looking at KEGG pathways using enrichr ^38,39^ in order to compare results.

## Results

There is limited evidence of pairwise genetic correlation among the four diseases using cross-trait LD score regression. The lack of an AD-PD pairwise correlation has already been reported, as well as between AD-MS and PD-MS.^40^ We also found no statistically significant evidence for genetic correlation between ALS-AD (0.2, p=0.08), ALS-PD (-0.08, p=0.01), and ALS-MS (-0.04, p=0.7).

For the grouped cell-type analysis from Finucane *et al*.,^31^ the most significant enrichment was seen for the immune/hematopoietic category for MS (10.1, p= 3.8 × 10^−13^), confirming the recognized role of the immune system in this disease. This category was also significantly enriched for heritability of AD (5.5, p= 2.4 × 10^−7^), in addition to liver (10.5, p= 1.1 × 10^−5^), and these AD signals remained significant even after the removal of APOE (chr19: 44905754-44909393) (5.5, p= 2.5 × 10^−7^ and 10.5, p= 1.1 × 10^−5^ respectively). For ALS and PD, there were no significantly enriched functional categories (Fig 1).

**Fig. 1.**
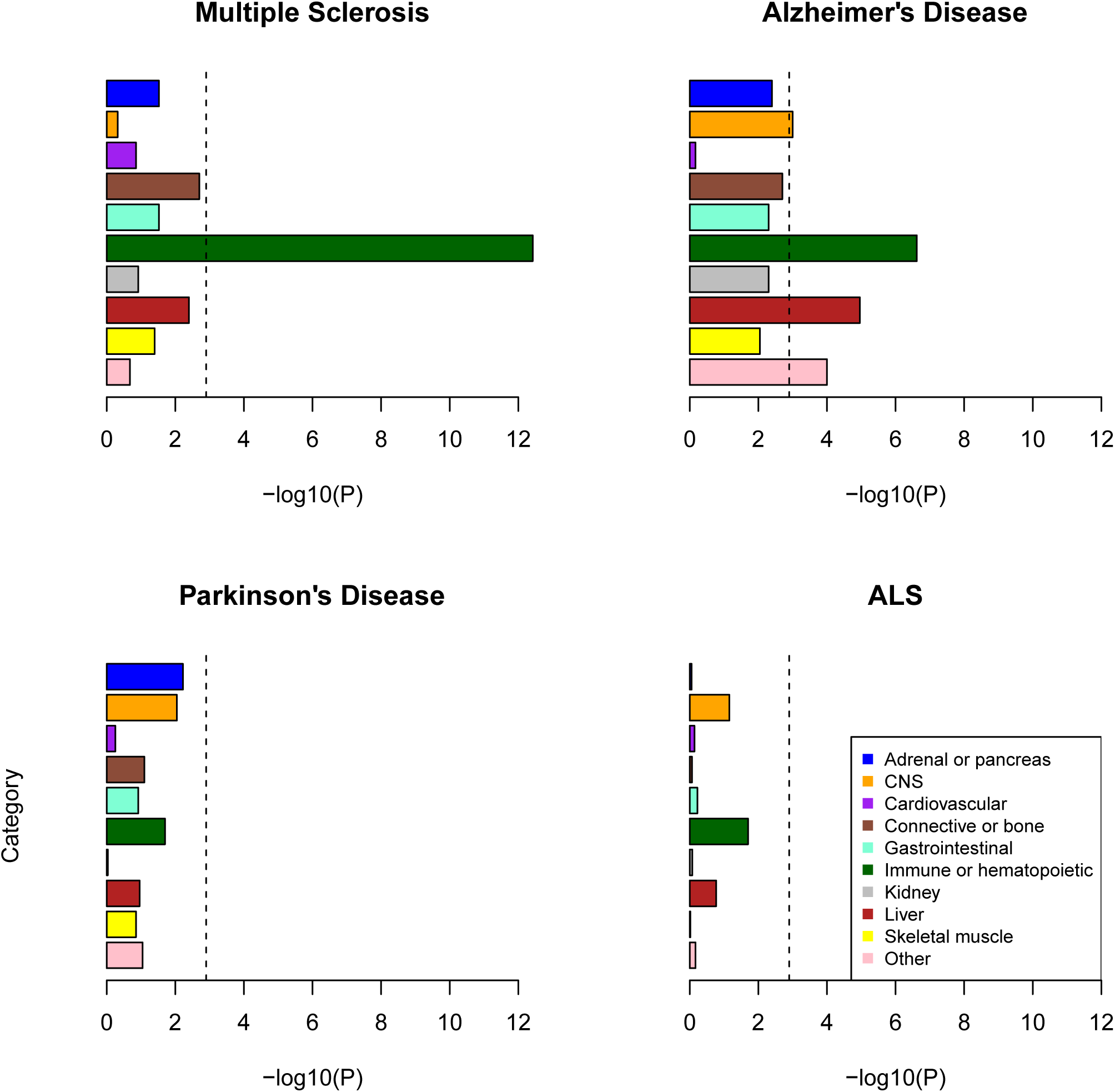
Enrichment of cell-type groups as used in Finucane et al. 2015. The black dashed lines at -log_10_(P) = 2.9 is the cutoff for Bonferroni significance.

At the tissue level, none of the enrichments were significant for the brain annotations. The most suggestive signal was for the inferior temporal region in AD (4.9, p= 6.6 × 10^−4^).

For the cell-specific immune annotations assessed relating to both the innate and adaptive immune systems, there was significant enrichment for MS heritability and to a lesser extent for AD and PD. There was no enrichment of heritability for ALS, the smallest dataset in our study (**Supplementary Table 1**, Fig 2). Strong MS signals for heritability enrichment were found in all immune cell categories, including both adaptive and innate cell types. Significant AD signals were found in all immune cell categories except for the non-T-cell/non-B-cell component of the adaptive immune system. For PD, only two annotations passed the multiple testing threshold: primary T helper cells PMA-I stimulated and primary T regulatory cells from peripheral blood (5.2 and 5.4, respectively, p= 0.0002 for both), but several other immune annotations were suggestive.

**Fig. 2.**
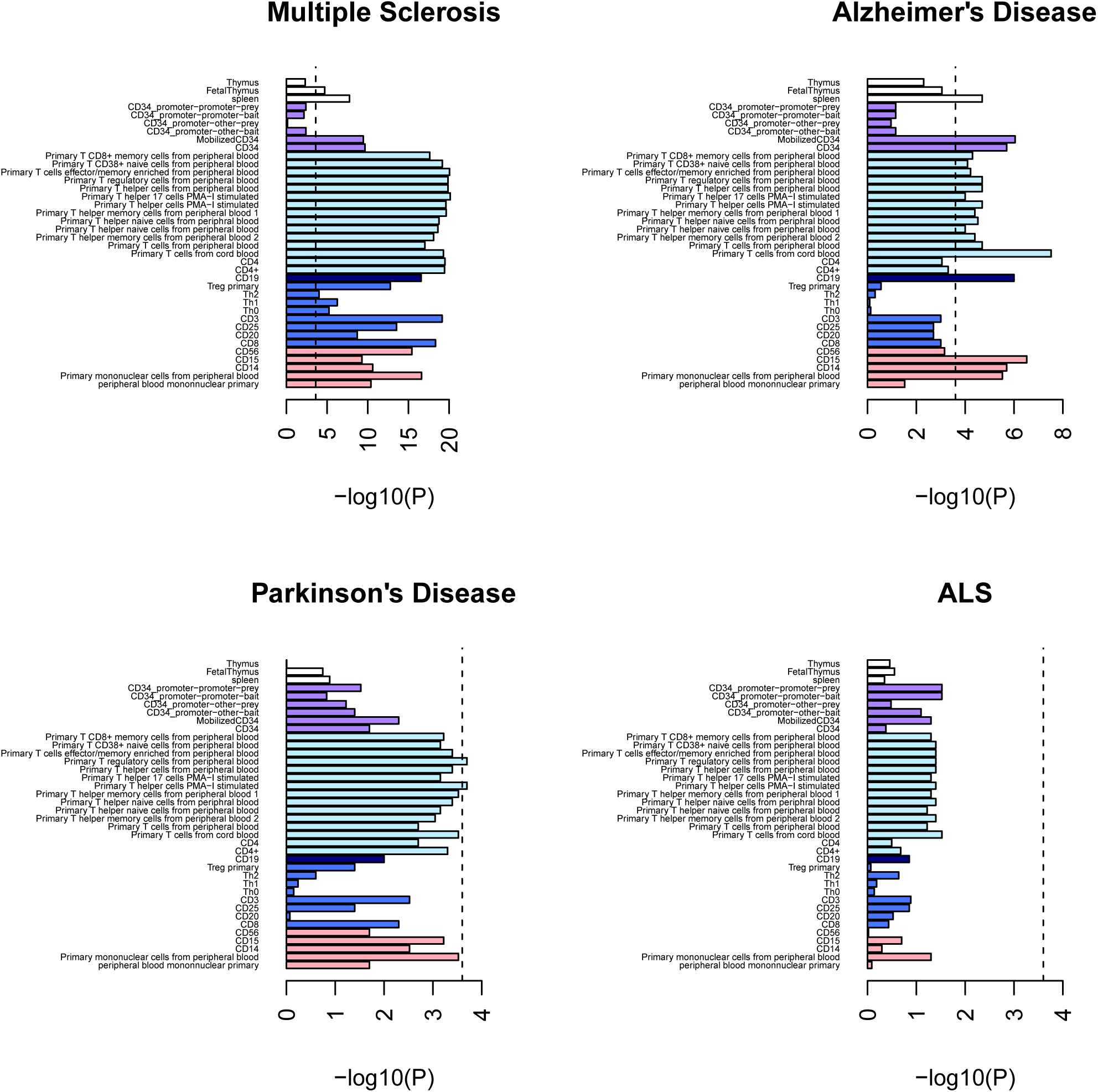
Enrichment of immune cell annotations. The black dashed lines at −log_10_(P) = 3.6 is the cutoff for Bonferroni significance. White bars= tissue; purple bars= CD34 (marker of immature hematopoietic cells – not strictly adaptive or innate); light blue bars= marker of T cells; dark blue bar= marker of B cells; royal blue bars= cells of the adaptive immune system; pink bars= cells of the innate immune system

Consistent with previous applications of the LD score regression method, we included the annotations separately in the regression model. This means that that enrichments in innate immune cells could in principle be due to overlap in annotation with adaptive immune cell-types, and vice versa. To assess this issue, we determined the degree of annotation overlap between all pairs of immune cell types in our study (**Supplementary Table 2**). We found the degree of overlap between innate versus adaptive cells ranged from 0.06% (for CD14: a marker for monocytes versus CD20: a marker of B lymphocytes) to 12% (for peripheral blood mononuclear primary cell versus primary T cells from cord blood), suggesting a large degree of independence between adaptive and innate cell marks. To further investigate this issue, we carried out deeper analyses on a representative adaptive cell type (primary T cells from cord blood) and a representative innate cell type (CD15:a marker for neutrophils), both of which displayed strong heritability enrichment signals in AD. The annotation overlap between these two cell types was 6.7% (Supplementary Table 2). When we included both annotations simultaneously in the LD score regression model, we found that both cell lines remained significantly enriched for MS (22.0, p= 7.4 × 10^−20^ and 17.2, 2.3 × 10^−5^, respectively). Similarly, for AD, both cell lines remained significantly enriched (8.7, p= 1.7 × 10^−7^ and 14.3, p= 7.3 × 10^−6^, respectively). Neither of these cell lines had reached significance for PD or ALS in the models where they were inputted separately, nor were they significant when included simultaneously into the model. The enrichment results for primary T cells from cord blood and CD15 when included simultaneously in the model for PD are 4.9, p= 1.1 × 10^−3^ and 7.0, p= 5.2 × 10^−3^, respectively; and for ALS 3.5, p= 0.03 and 3.1, p= 0.37 respectively. Overall, these analyses provided us with re-assurance that we were detecting independent signals in adaptive versus innate immune cell types.

Our heritability enrichment analysis within brain-related and immune-related gene sets also provided strong evidence for a signal in the immune gene set, and not in the brain gene set (**Supplementary Table 1**). As expected, the strongest immune gene signal was for MS (1.6, p= 4.6×10^−14^). We have previously reported the enrichment of this immune gene list in the same MS dataset, using an earlier version of LD score regression.^32^ The immune gene list was also enriched for heritability in AD (5.2, p= 4.8 × 10^−4^), and the effects in PD and ALS were suggestive but would not survive multiple testing correction (4.5, p= 0.02 and 2.5, p= 0.03, respectively). The brain gene list was not significantly enriched in any of the neurodegenerative diseases assessed (among the other three diseases enrichment ranges from 0.9 to 1.9, p >0.04 for all three).

Finally, we compared the above results to an Ingenuity IPA pathway enrichment analysis, both within canonical pathways (**Supplementary Table 3a**) and within diseases/biological functions including cancer-related functions (**Supplementary Table 3b**). We also compared our results to an enrichr pathway enrichment analysis (**Supplementary Table 3c**). Remarkably, for the IPA canonical pathway analysis, all the significant pathways save one (“Aldosterone Signaling in Epithelial Cells”) were found to be connected to either adaptive or innate immune response. Specific examples included: in MS (e.g. T helper cell differentiation, role of macrophages, fibroblasts and endothelial cells in RA, B cell receptor signaling, dendritic cell maturation, PI3K signaling in B lymphocytes, CD40 signaling; PKCθ signaling in T lymphocytes, NF-κB activation by viruses); in PD (e.g. dendritic cell maturation – shared with MS, graft-versus-host disease signaling, altered T cell and B cell signaling in rheumatoid arthritis); in AD (IL-8 signaling, IL-12 signaling and production in macrophages, Fc epsilon RI signaling, Fc*γ* receptor-mediated phagocytosis in macrophages and monocytes, role of pattern recognition receptors in recognition of bacteria and viruses, natural killer cell signaling); and in ALS (e.g. NF-κB signaling). The value of the IPA method was also demonstrated in providing significant signals for other pathways previously implicated in the pathogenesis of AD, including CREB signaling in neurons,^41^ neuregulin signaling, and ErbB signaling.^42^ For the IPA diseases/biological functions analysis, various cancers came up as most strongly significant for all the disorders. Cancer has been shown to be correlated with multiple immune disorders ^43^, and there is evidence of cancer and neurodegenerative disorders, such as PD, sharing common pathways.^44^ The enrichr analysis revealed many significant immune-related pathways, in line with the IPA canonical pathways analysis.

## Discussion

Multiple lines of evidence suggest a significant contribution of variants exhibiting functional marks for chromatin accessibility (i.e. histone marks, DNase I hypersensitive sites) in immune cell types to the heritability of two neurodegenerative diseases, namely AD and PD. Annotations from immune cells are most significantly enriched for the heritability of MS, a known autoimmune disease which acted as a positive control in our investigations.^17^ Immune annotations are also consistently enriched but to a lesser degree for AD (with involvement from both the innate and adaptive immune systems), and some cell-specific immune annotations (T-cells) were significantly enriched for PD. A lack of results from the ALS dataset could be attributed to this dataset being smaller than the other datasets investigated (Table 1). These results provide further support for the neuroinflammatory hypothesis of neurodegenerative disease,^15,16^ and highlight the potential utility of immune modulating agents, such as those currently used in MS for the treatment of AD and PD. However, one needs to be cautious with interpreting these cell/tissue-type specific results in the absence of functional and other studies.

We note that if we correct for the 17 GWASs assessed in Finucane *et al*.^31^ as well as the four GWASs we assessed here for the 10 cell-type groups ((17+4) × 10 = 210 hypotheses tested), both the immune/hematopoietic and liver categories remain significant for AD.

The role of the immune system in AD pathogenicity has been previously shown^22,25^ and previous pathway analysis of the AD GWAS we assessed here showed enrichment in immune-related pathways.^24^ Findings are strongest for the innate immune response, for instance association with the *TREM2* gene, which in brain cells are primarily expressed on microglia.^45,46^ Our findings further support the role of immune variation in AD susceptibility. Interestingly, using LD score regression, AD was found to be not significantly correlated with a variety of immune diseases.^34^ This lack of correlation could be because when considering the entire genome the signal coming from the correlated loci between the diseases is diluted, or the immune variants involved in AD are different from those involved in other immune diseases. Microglia, the main immune cell type in the brain, have a different developmental trajectory separate from the peripheral immune system.^47^ The unique mechanisms of immune surveillance in the brain ^48,49^also makes immune diseases of the brain biologically distinct to peripheral immune diseases, but there is much evidence that disruption of the brain’s immune surveillance is critical to the “vicious cycle” of worsening pathology seen in neurodegeneration.^50^ Our analysis suggests a predominantly epigenomic mechanism for immune dysregulation in neurodegenerative disease, and if confirmed this may be of therapeutic relevance, as many drugs are known to act through this mechanism. Some, such as histone deacetylase inhibitors, could potentially be efficacious in neurodegenerative diseases.^51^

Functional marks from liver were also enriched for the heritability of AD. This result agrees with findings in the literature of the contribution of lipid metabolism through liver X receptors (LXR) to the initiation and progression of this disease.^52,53^

Canonical pathway analysis showed enrichment of AD associations in CREB signaling in neurons, and also IL8 and IL12 signaling (which are CREB regulated), supporting the immune hypothesis in AD, and pointing to interleukin signaling as a potential CREB-responsive mechanism.^54^

Our results do not provide statistically significant evidence that variants overlapping with functional annotations from the brain contribute excessively to the heritability of neurodegenerative diseases. The brain doubtless plays an important role in the genetic aetiology of these diseases. The lack of brain annotation enrichment could be due to data being based on few samples for the brain. Furthermore, the brain is a very heterogeneous tissue. Data from brain regions contain a mixture of different cell types such as microglia and neurons. Single cell sampling may reduce this heterogeneity in the future.^55^ This analysis should be revisited as brain annotation information improves.

In summary, our results suggest a significant contribution of variants that exhibit chromatin accessibility marks in immune cells to the heritability of two neurodegenerative diseases, namely AD and PD.

## Acknowledgements

We would like to thank Andy Singleton and Mike Nalls for access to the Parkinson’s disease summary statistics. We would also like to thank Isabella Fogh and John Powell for providing access to the ALS summary statistics. The SLAGEN (Italian Consortium for the Genetics of ALS) and ALSGEN (International Consortium of Amyotrophic Lateral Sclerosis Genetics) Consortia contributed ALS/controls samples. The multiple sclerosis summary statistics were accessed through Immunobase (https://www.immunobase.org/). We thank Borbala Mifsud for advice on HiC datasets. We also thank Jayne Danska for input on the immune interpretation.

Funding for this work: SAG was funded through the Weston Brain Institute International Fellowship in Neuroscience. JGP was funded by Fulbright Canada, Weston Canada, and Brain Canada through the Canada Brain Research Fund, a public-private partnership established by the Government of Canada. MRB was funded through the National Institutes for Health Research (NIHR) as part of the portfolio of translational research of the NIHR Biomedical Research Unit at Barts. JH, MR and MEW were funded through Medical Research Council (MRC) grant number G0901254/1.

We thank the International Genomics of Alzheimer’s Project (IGAP) for providing summary results data for these analyses. The investigators within IGAP contributed to the design and implementation of IGAP and/or provided data but did not participate in analysis or writing of this report. IGAP was made possible by the generous participation of the control subjects, the patients, and their families. The i–Select chips was funded by the French National Foundation on Alzheimer’s disease and related diseases. EADI was supported by the LABEX (laboratory of excellence program investment for the future) DISTALZ grant, Inserm, Institut Pasteur de Lille, Université de Lille 2 and the Lille University Hospital. GERAD was supported by the Medical Research Council (Grant n° 503480), Alzheimer’s Research UK (Grant n° 503176), the Wellcome Trust (Grant n° 082604/2/07/Z) and German Federal Ministry of Education and Research (BMBF): Competence Network Dementia (CND) grant n° 01GI0102, 01GI0711, 01GI0420. CHARGE was partly supported by the NIH/NIA grant R01 AG033193 and the NIA AG081220 and AGES contract N01–AG–12100, the NHLBI grant R01 HL105756, the Icelandic Heart Association, and the Erasmus Medical Center and Erasmus University. ADGC was supported by the NIH/NIA grants: U01 AG032984, U24 AG021886, U01 AG016976, and the Alzheimer’s Association grant ADGC–10–196728.

## Author Contributions

S.A.G., M.R., M.R.B., J.K., and M.E.W. are responsible for concept and study design. S.A.G. analysed data, M.R.B. conducted the pathway analysis, and J.G.P. analysed data for the gene list enrichment. S.A.G. drafted the manuscript and figures, with input from all the authors.

## Supplementary files

**Supplementary Table 1**. Annotation enrichment results. Red cells mark enrichment that survived Bonferroni correction.

**Supplementary Table 2**. Overlap among chromatin accessibility annotations for immune cells. The main diagonal shows genome coverage (base pairs) for that cell type. The upper off-diagonal shows the overlap coverage (base pairs) for that cell-type-pair. The lower off-diagonal shows the proportion of overlap coverage for that cell-type pair.

**Supplementary Table 3**. **[a]** Ingenuity Pathway Analysis (IPA) results for canonical pathways. Red cells mark enrichment that survived Bonferroni correction. **[b]** Ingenuity Pathway Analysis (IPA) results for cancer-related functions. Red cells mark enrichment that survived Bonferroni correction. **[c]** enrichr KEGG pathway results. Multiplication of the p-value computed using the Fisher exact test with the z-score of the deviation from the expected rank as described in the enrichr paper.^38^

